# XTalkiiS: a tool for finding data-driven cross-talks between intra-/inter-species pathways

**DOI:** 10.1101/437541

**Authors:** A. K. M. Azad

## Abstract

Cell-cell communication via pathway cross-talks within a single species have been studied *in silico* recently to decipher various disease phenotype. However, computational prediction of pathway cross-talks among multiple species in a data-driven manner is yet to be explored. In this article, I present XTalkiiS (Cross-talks between inter-/intra species pathways), a tool to automatically predict pathway cross-talks from data-driven models of pathway network, both within the same organism (intra-species) and between two organisms (inter-species). XTalkiiS starts with retrieving and listing up-to-date pathway information in all the species available in KEGG database using RESTful APIs (exploiting KEGG web services) and an in-house built web crawler. I hypothesize that data-driven network models can be built by simultaneously quantifying co-expression of pathway components (i.e. genes/proteins) in matched samples in multiple organisms. Next, XTalkiiS loads a data-driven pathway network and applies a novel cross-talk modelling approach to determine interactions among known KEGG pathways in selected organisms. The potentials of XTalkiiS are huge as it paves the way of finding novel insights into mechanisms how pathways from two species (ideally host-parasite) may interact that may contribute to the various phenotype of interests such as malaria disease. XTalkiiS is made open sourced at https://github.com/Akmazad/XTalkiiS and its binary files are freely available for downloading from https://sourceforge.net/projects/xtalkiis/.

## Introduction

In biological systems, genes doesn’t work alone; rather as a group they perform certain activities named as pathways. Sometimes, the activities of certain pathways even gets modified by others pathways due to the interference among them, a phenomenon called ‘pathway cross-talks’. In many cases such interference plays critical roles in mediating novel mechanisms of certain diseases-associated activities e.g. tumor progression or acquired drug resistance. Moreover, these cross-talks may also be present not only within the same organism but also between organisms e.g. interference between host-parasite pathway activities.

Development of many complex diseases such as cancer often happens due to the genetic/epigenetic alteration of some key driver genes and their perturbed influence through their pathway activities [5. Moreover, recent researches *in vivo* and *in vitro* have shown that cancer cells acquires resistance to particular inhibitors by adapting their signalling circuitry, activation of alternate pathways and cross-talks among various signalling pathways [5]. Again, Komurov *et al.* reported that regulatory pathways such as *glucose deprivation response pathway* cross-talks with *EGFR*-mediated pathways to provide cancer cells an alternate route for glucose intake, which is essential for their survival for which tumor ultimately relapses [9]. So, for better understanding of pathway activities within a biological systems under various research hypotheses, it is often crucial to study the pathway cross-talks within the organism.

Moreover, it is important to study interactions among multiple organisms, especially within microbial ecosystems [10] for several reasons: 1) how their pathways are interconnected, 2) how their interactions changes due the perturbation of one or both of their systems, 3) how do they correspond to external factors such environmental changes or treatment stimuli, etc. Hence, more research required to determine and characterize these inter-species pathway cross-talks to reveal better insights into the networks of diseases-associated biological pathways in a data-driven manner.

Gene or protein co-expression measurements using high-throughput data sets can model their relationships/dependencies in performing a particular biological activities [6] in a particular pathway in a data-driven manner. These putative relationships can form a network structure mimicking pathways [6], and their interactions in a data-driven manner [5], where the nodes are individual genes/proteins and edges are the relationships among them within that particular species. I hypothesize that similar approach could be applied to study the relationships among pathway components from multiple species by forming a mega network structure, which can be determined by, 1) first measuring genome-wide high-throughput information of pathway genes/proteins simultaneously in multiple organism, and then 2) evaluating their co-expression.

Data-driven models of pathway network provide useful ways of capturing dynamic patterns in pathway activities in a context-specific manner [2]. However, typical definitions of pathway cross-talks aren’t suitable for data-driven models of pathway networks as they may have novel dependencies among genes/proteins within those network structure [2]. Hence, we defined a novel cross-talk categorization namely *Type-I* and *Type-II* cross-talks that are suitable for data-driven network models and covers all the cross-talk definitions from the state-of-the-art approaches [2].

In this study, we proposed a framework and a software tool called XTalkiiS (Cross-talks between inter-/intra species pathways) that uses our previously proposed cross-talk modeling suitable for data-driven pathway networks [2] and determine pathway cross-talks within the same species (intra-species) and between two-component biological systems i.e. host-parasite manner.

## Implementation

Figure 1 demonstrates the main interface of XTalkiiS. XTalkiiS operates to find pathway cross-talks within the same species (or organism) or between *two* species. Upon loading the main interface, XTalkiiS retrieves all the organism names from KEGG database by making HTTP web request via a RESTful API call, and lists them in two combo-boxes side-by-side from where users can select two organisms. Upon selecting an organism from each of these two combo-boxes, all pathways names available in the KEGG databases for corresponding organisms are again retrieved using another RESTful API call. When all the pathways are listed for those two organisms, some (or all) of them can be selected which populate two corresponding list-boxes. Next, user loads a gene-gene dependency file from local machine which has information of following structure,

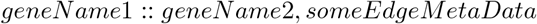

where *geneName*1 and *geneName*2 belongs to *pathway*1 and *pathway*2, respectively which could be from the same organism (if the same organism is selected in both of the combo-boxes) or two different organisms. This gene-gene dependency file saves the adjacency information of various pathway components in the mega network structure composed of various pathways from the same or different species.

**Figure 1.**
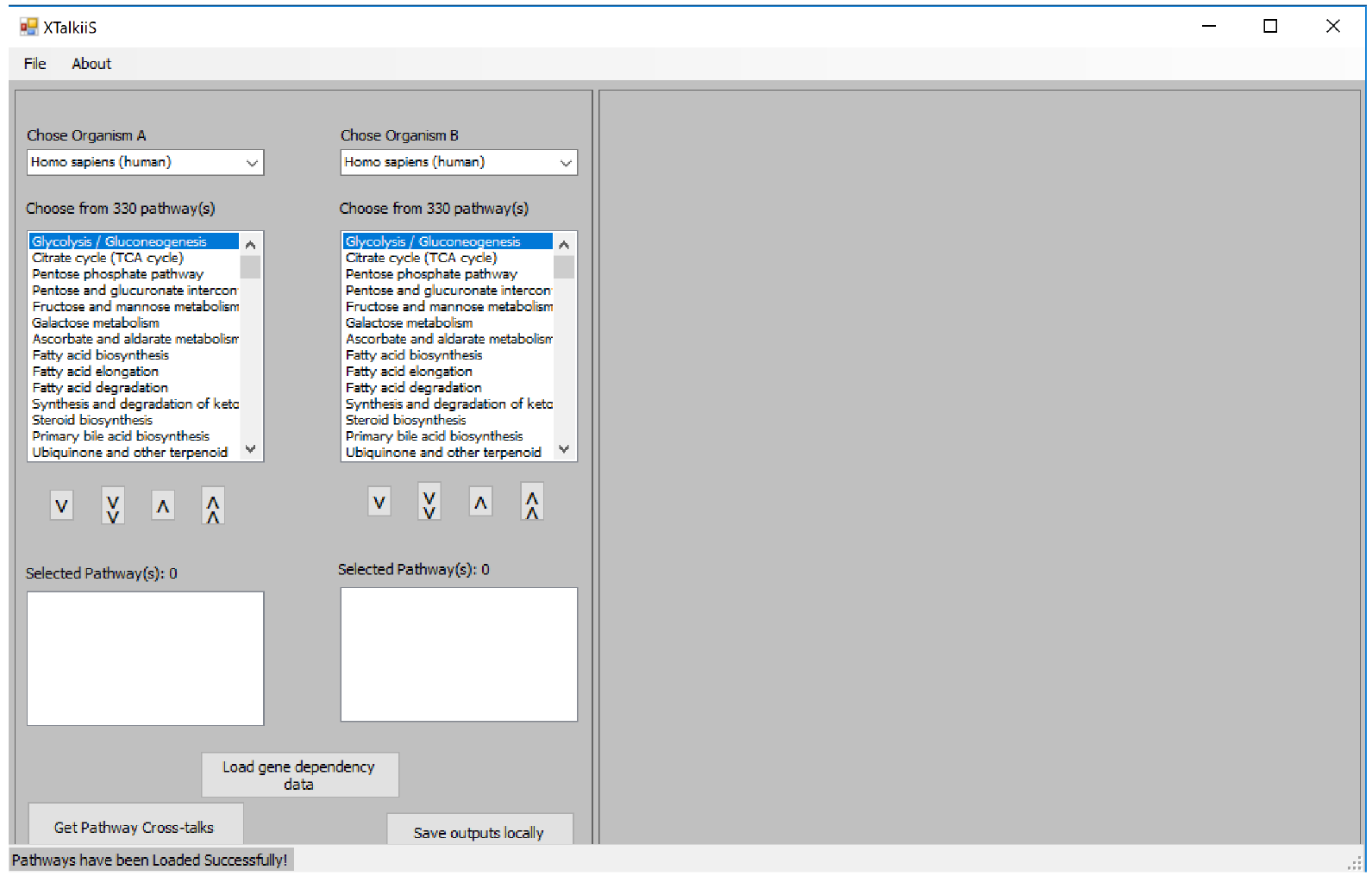
Main interface of XTalkiiS tool

Then upon pressing a button it will retrieve all the up-to-date gene names in all the pathways in those two list-boxes via another RESTful API call, which is similar to a previously developed tool called KPGminer [4]. These API calling protocols are listed in the KEGG website for developers’ use. In the same event, it applies the cross-talk categorizations [2] on the gene-gene dependency file loaded before and finds two types of cross-talks, *Type-I* and *Type-II* cross-talks. Next, the output panel on the right shows 1), all the pathway genes in the first grid-view, and 2) all the *Type-I* and *Type-II* cross-talks among all the pathways in intra-/inter-species manner in the second grid-view.

**Table 1.**
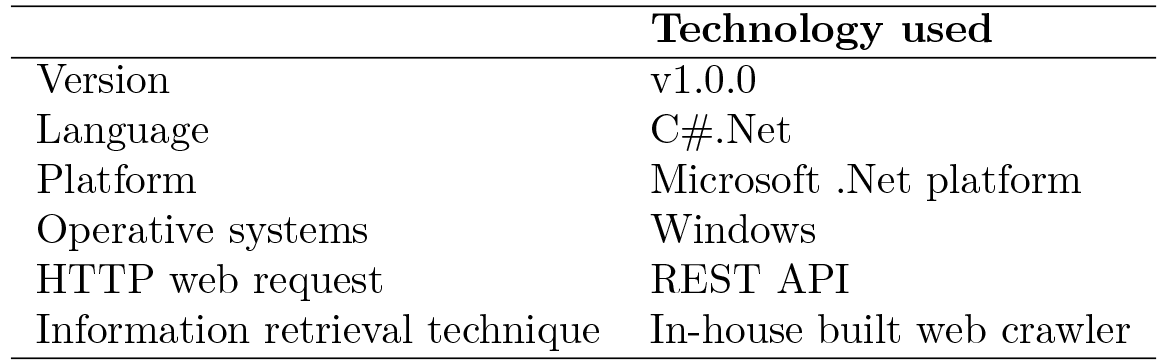
XTalkiiS Metadata

|  | Technology used |
| --- | --- |
| Version | v1.0.0 |
| Language | C#.Net |
| Platform | Microsoft .Net platform |
| Operative systems | Windows |
| HTTP web request | REST API |
| Information retrieval technique | In-house built web crawler |

## Application: Gene expression data from GSE16179

Next, I apply XTalkiiS on gene-gene dependency data set collected from [3], which is a list of aberrant gene-pairs derived from lapatinib-sensitive and lapatinib-resistant BT474 cell-lines [GSE16179] using Bayesian statistical modelling [for details see Methods section from [3]]. This example demonstrates the data-driven pathway cross-talks from human breast cancer samples, hence indicating intra-species pathway cross-talks. This network consists of 76,476 aberrant gene-pairs with corresponding edge label as edge meta-data.

**Figure 2.**
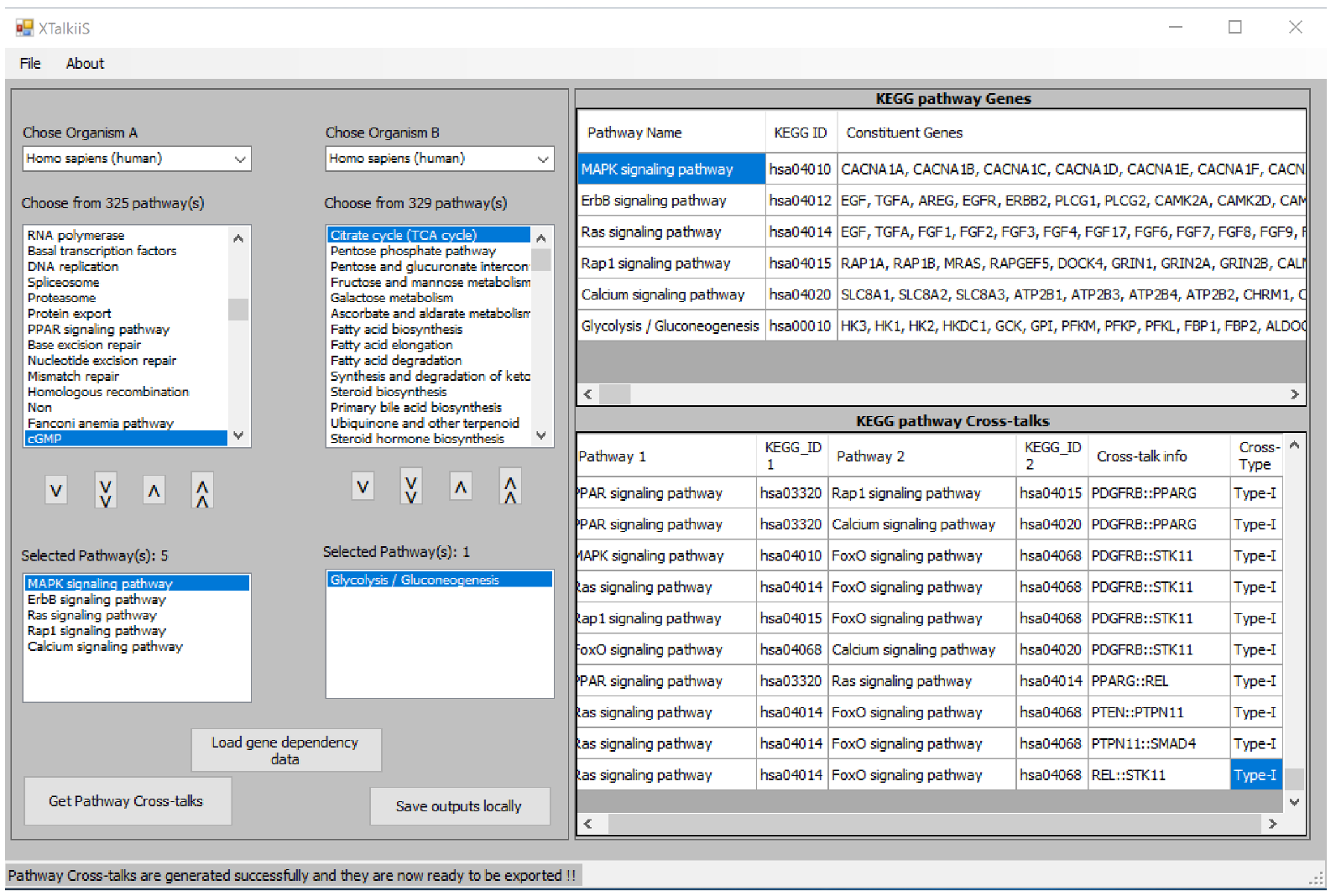
XTalkiiS detects pathway cross-talks from GSE16179 data

Figure 2 shows the output of XTalkiiS after running with the above dataset. It retrieves pathways from MAPK signalling, ErbB signalling, Ras signalling, Rap1 signalling, Calcium signalling, and Glycolysis/Gluconeogenesis pathways from Homo Sapiens and finds cross-talks among them.

## Discussion

Biological systems often reveals interactions of multiple pathways as various cell-cell communications. Small organisms may reside in host organism and their pathways may interact with the pathway in host organisms in order to express certain disease phenotype in a host-parasite manner such as malaria disease. In this regard, XTalkiiS has tremendous potential in finding these inter-species pathway cross-talks as it may decipher novel mechanisms of various biological activities in various research hypotheses in a data-driven manner.

This version of XTalkiiS has one limitation: it loads pathway genes for each individual pathways via making HTTP web request iteratively for each pathways which may be time-consuming depending on the number of pathways. In the next versions, it will adapt multi-threading approach where each thread will make separate HTTP web request, especially because the gene loading step for each pathways are independent of each others. Moreover, XTalkiiS can be resourced with more pathway databases including Reactome [7], Wikipathways [8] or GO terms [1]. I hope, XTalkiiS will be a resourceful tool in future, when finding inter-species pathway cross-talks will become a necessary research component in the field of computational biology.

